# Tropism of AAV.CPP.16 in the respiratory tract and its application for a CRISPR-based gene therapy against SARS-CoV-2

**DOI:** 10.1101/2023.11.17.567583

**Authors:** Zhi Yang, Yizheng Yao, Xi Chen, Victoria Madigan, Xianqun Fan, Jun Pu, Fengfeng Bei

**Affiliations:** Department of Neurosurgery, Brigham and Women’s Hospital, Harvard Medical School, Boston, MA 02115, USA; Department of Ophthalmology, Ninth People’s Hospital, Shanghai JiaoTong University School of Medicine, Shanghai 200011, China; Jiangsu Key Laboratory of Neuropsychiatric Diseases and Institute of Neuroscience, Soochow University; Clinical Research Center of Neurological Disease, The Second Affiliated Hospital of Soochow University, Suzhou 215123, China; Department of Neurosurgery, The Second Affiliated Hospital of Kunming Medical University, Kunming 650106, China; NHC Key Laboratory of Drug Addiction Medicine, Kunming Medical University, Kunming 650500, China; Broad Institute of MIT and Harvard, Cambridge, MA 02142, USA; Department of Biological Engineering, Massachusetts Institute of Technology, Cambridge, MA 02139, USA

**Keywords:** AAV, lung, CRISPR, Cas13d, SARS-CoV-2

## Abstract

Efficient gene delivery vectors are essential for developing gene therapies for respiratory diseases. Here, we report that AAV.CPP.16, a novel AAV9-derived adeno-associated virus vector, can efficiently transduce airway epithelium systems and lung parenchyma cells in both mice and non-human primates after intranasal administration. AAV.CPP.16 outperforms AAV6 and AAV9, two wild-type AAVs with demonstrated tropism to respiratory tract tissues, and can target major cell types in the respiratory tract and the lung. We also report an “all-in-one”, CRISPR-Cas13d-based AAV gene therapy vector that targets the highly conserved RNA-dependent RNA polymerase (Rdrp) gene in SARS-CoV-2, and show the potential of such gene therapy against a broad range of circulating and emergent SARS-CoV-2 variants. Thus, AAV.CPP.16 could be a useful gene delivery vector for treating genetic respiratory diseases and airborne infections including for developing a potential prophilaxis to SARS-CoV-2.

## Introduction

Progress in respiratory gene therapy and gene editing is hampered by a lack of efficient delivery vectors. Adeno-associated virus (AAV) is a commonly used viral vector for *in vivo* gene delivery with accumulating clinical trial experiences^1^. Among the wild-type AAVs, AAV6 and AAV9 are two promising serotypes for targeting the respiratory tract and the lung^2^. For example, AAV9 has been shown to transduce mouse airway and lung tissues after intranasal administration; it can be readministered as early as one month after initial application and can be used for intranasal antibody gene therapy against pandemic influenza in animal models^3–5^. Recently, we reported an AAV9-derived capsid AAV.CPP.16 with species-conserved neurotropism after systemic administration^6^. During characterization of vector biodistribution, we also observed enhanced transduction of the lung by AAV.CPP.16 relative to its parent capsid AAV9. Thus, we decided to further explore the use of AAV.CPP.16 as an intranasal gene transfer vector for gene therapy and gene editing for potential treatment of respiratory genetic diseases and airborne viral infection.

Vaccination against SARS-CoV-2 has been highly effective in combating the COVID-19 pandemic^7^. However, such preventiave strategy is less effective or even invalid in vulnerable populations of immune-deficient patients^8^. Furthermore, constant mutation and evolution of SARS-CoV-2 viruses have led to the emerging of SARS-CoV-2 variants that could potentially break through the protective barrier established by active vaccination and/or passive viral exposure^9^. Continuous research on non-immune-based antiviral approaches may provide additional prevention and treatment options for SARS-CoV-2 infection^10^.

In this study, we systemically characterize AAV.CPP.16-mediated gene delivery for major cell types that belong to the respiratory tract using cell culture, mouse and non-human primate models. In addition, we design and test a new, gene therapy-based, passive prophylaxis to SARS-CoV-2 injection using a CRISPR (Clustered Regularly Interspaced Short Palindromic Repeats) system.

## Results

### Tropism of AAV.CPP.16 for airway cells and mouse respiratory tract

We first examined the transduction efficiency of AAV.CPP.16 in cultured airway cells and in mice and compared with AAV6 and AAV9, two previously reported respiratory-tract-targeting AAVs ^11–15^ (**Fig. 1a**). Viral vectors were packaged carrying a GFP reporter with a ubiquitous CAG promoter (ssAAV-CAG-GFP). RPMI 2650 cell, a human nasal epithelial cell line, was prepared and treated with AAVs for 4 days using a multiplicity of injection (MOI) of 1 × 10^5^. It was found that the average fluorescence intensity of GFP expressed by AAV.CPP.16 was 3.0-fold and 6.6-fold greater than AAV6 and AAV9 respectively (P = 0.0023, P = 0.0006 respectively) (**Fig. 1b,c**). No significant difference of GFP signals between AAV6 and AAV9 was observed.

**Fig. 1.**
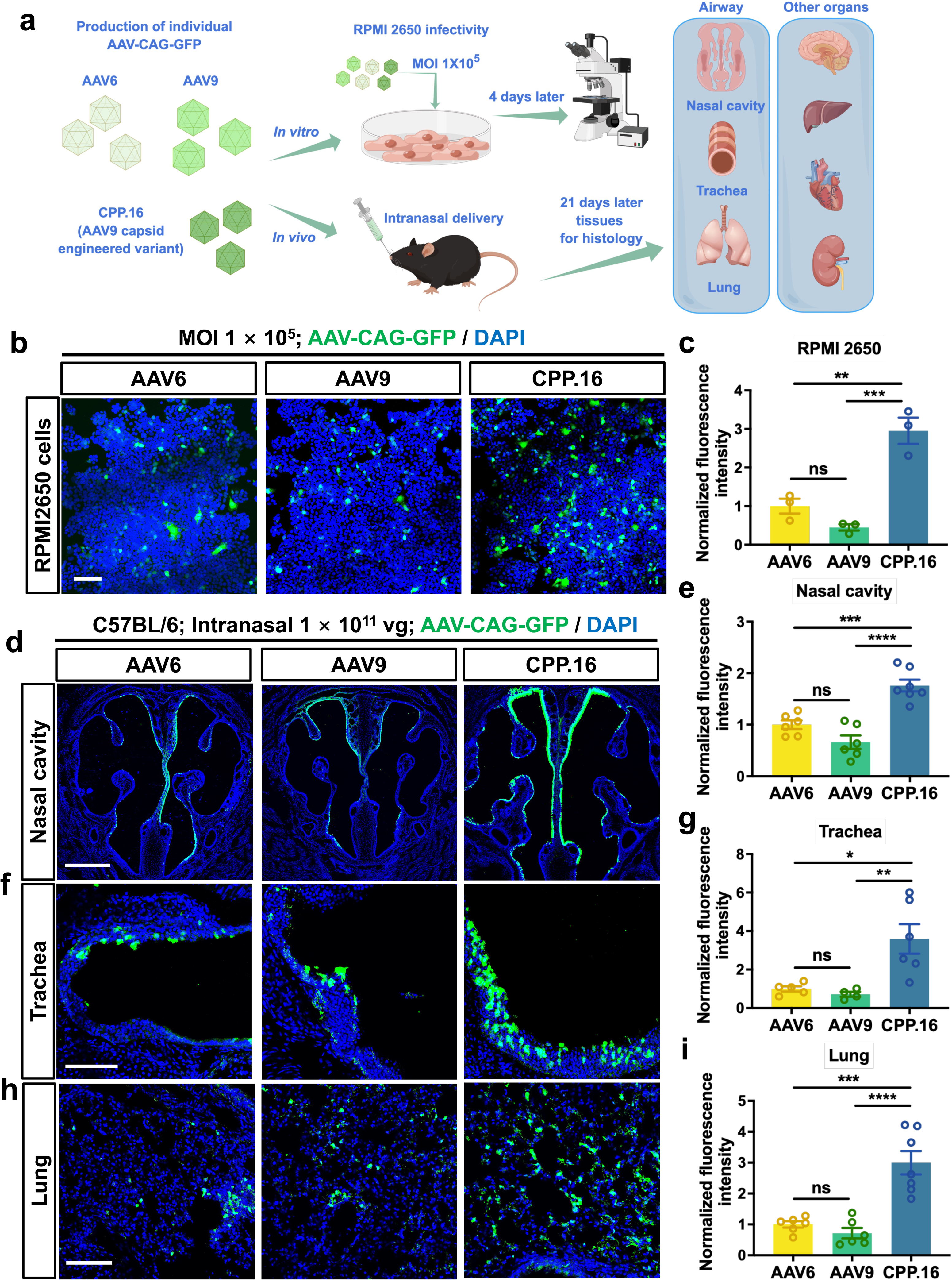
AAV transduction of airway cells in cell culture and mouse models. **a,** Schematic diagram showing experimental design. **b, c,** Images showing transduction of cultured human nasal epithelial cells RPMI 2650 in vitro (Multiplicity of Infection, MOI, 1 × 10^5^) by AAV6-CAG-GFP, AAV9-CAG-GFP, and AAV.CPP.16-CAG-GFP (CPP.16) (b) and quantification of GFP fluorescence intensity normalized as fold-change relative to AAV6 (c). Mean ± s.e.m., n = 3. Scale bar: 100 μm. **d-I,** Representative fluorescence images and quantitative analysis showing AAV-mediated GFP expression in the nasal cavity **(d, e)**, trachea **(f, g)**, lung **(h, i)** in C57BL/6J mice 21 days after intranasal delivery of 1 × 10^11^ vg of AAV6, AAV9 or CPP.16. Mean ± s.e.m., n = 6-7 each treatment. Scale bar in **d**: 500 μm. Scale bar in **f, h**: 100 μm. Statistical analyses were performed using one-way ANOVA with Tukey’s multiple comparison test (\**p* < 0.05, \*\**p* < 0.01, \*\*\**p* < 0.001, \*\*\*\**p* < 0.0001).

To test AAV.CPP.16 in mice, intranasal administration of 1 × 10^11^ vg AAVs-CAG-GFP per animal was performed using both nostrils in C57BL/6J mice. 3 weeks later, nasal cavity, trachea and lung tissues were fixed and harvested for assessing AAV-mediated GFP expression, while other organs were collected for further biodistribution analysis. The fluorescence images revealed superior fluorescence intensity in the nasal cavity of mice treated with AAV.CPP.16 (2.7-fold vs. AAV9 and 1.8-fold vs. AAV6, **Fig. 1d,e**). Strong and broad GFP signal was observed in membraneous structures in anatomical regions including the nasal septum, nasal turbinates and different regions of nasal meatuses, while only partial and weak expression of GFP was noticed in nostrils treated with AAV6 or AAV9 (**Fig. 1d**). No GFP expression was observed in deep tissues in the nasal cavity. A similar examination revealed significantly enhanced transduction efficiency by AAV.CPP.16 in inner linings of the trachea (**Fig. 1f,g**) and the lung (**Fig. 1h,i**) as compared with AAV6 and AAV9. No statistical differences were observed when comparing AAV6 to AAV9 in the nasal cavity, trachea and lung tissues. In addition, intranasal injection of AAV6, AAV9, and AAV.CPP.16 resulted in no to little transduction in the brain, retina, tongue, liver, spleen, intestine, stomach, kidney and muscle (**Supplementary Fig. S1**), suggesting relative selectivity of intranasally administered AAVs for targeting the respiratory tract. A small amount of transduced cells were observed in the liver in mice treated with AAV.CPP.16.

Next, we examined the types of AAV-transduced airway cells using antibody markers (**Fig. 2a**). In the nasal cavity, immunostaining against MUC5A and α-Tubulin^16^ were used for labeling the two primary cell types, namely, the goblet- and ciliated cells respectively. We found that AAV.CPP.16-GFP of 1 × 10^11^ vg transduced approximately 42.7% of MUC5A-positive goblet cell, as compared with much lower transduction percentages of 3.0% for AAV6 and 7.8% for AAV9 (**Fig. 2b,e**). AAV.CPP.16-GFP also appears to transduce more α-Tubulin-positive ciliated cells per unit areas in the nasal cavity in comparison to AAV6-GFP and AAV9-GFP (**Supplementary Fig. S2a**), although the percentage transduction of ciliated cells could not be determined using α-Tubulin as a marker.

**Fig. 2.**
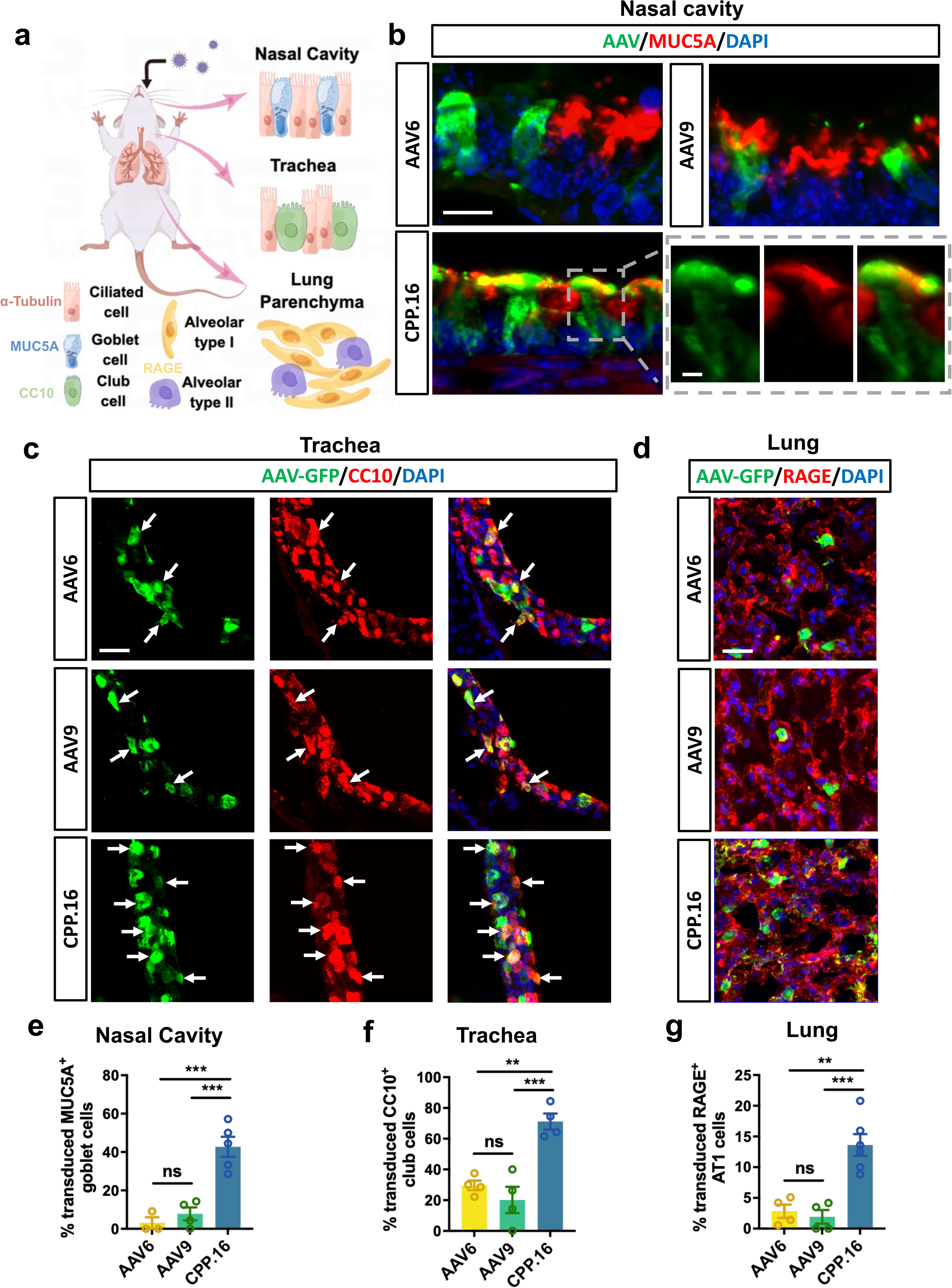
Characterization of AAV-transduced cell types in the mouse respiratory tract and the lung. **a,** Schematic diagram illustrating major airway cell types, their anatomical locations, and antibody markers. Mouse tissues were collected 21 days after intranasal administration of 1 × 10^11^ vg of AAV-CAG-GFP. **b,** Representative images showing transduced goblet cells (MUC5A^+^) in the nasal cavity. Scale bar: 25 μm for low-magnification images and 5 μm for enlarged images. **c,** AAV-mediated transduction of club cells (CC10^+^) in the trachea. Arrows point to cells co-labeled with GFP and CC10 antibodies. Scale bar: 25 μm. **d**, AAV-mediated transduction of alveolar type I (AT1) (RAGE^+^) in lung parenchyma tissues. Scale bar: 25 μm. **e-g**, Quantifications on the percentage of GFP^+^ goblet cells in the nasal cavity (e), club cells in the trachea (f) and AT1 cells in the lung (g). Mean ± s.e.m., n = 3-4 for each group. Statistical analyses were performed using one-way ANOVA with Tukey’s multiple comparison test (\*\**p* < 0.01, \*\*\**p* < 0.001).

In the trachea, we analysed the two major component cells, club- and ciliated cells, using antibody markers CC10^17^ and α-Tubulin respectively. AAV.CPP.16-GFP transduced 71.0% of the club cells, exhibiting significantly higher transduction efficiencies as compared with AAV6 (29.6%) and AAV9 (20.2%) (**Fig. 2c,f**). More GFP-positive cells were also observed with co-labeling for α-Tubulin in AAV.CPP.16-treated trachea tissues (**Supplementary Fig. S2b**). No significant difference in transduction was observed in either club or ciliated cells between AAV6 and AAV9 treatment.

Lastly, to characterize AAV transduction in the lung, we focused on measuring the AAV expression on alveolar type I (AT1) cells in lung parenchyma, which cover more than 95% of alveolar surface^18^, as well as ciliated cells in lung bronchi. We applied an antibody against RAGE^18^, a membrane protein, to determine AT1 cells. Intranasal administration of AAV.CPP.16 resulted in increased transduction of RAGE-positive AT1 cells (13.6% for AAV.CPP.16 vs. 2.8% for AAV6 and 1.9% for AAV9) (**Fig. 2d,g**). For ciliated cells located in the inner wall of lung bronchi, more of them were also transduced by AAV.CPP.16 as compared with AAV6 and AAV9 (**Supplementary Fig. S2c**),

### AAV.CPP.16 maintains superior tropism for the respiratory tract in NHPs

To test if the enhanced tissue tropism of AAV.CPP.16 in the respiratory tract can translate from mice to NHPs, we compare AAV.CPP.16 to AAV9 in adult cynomolgus macaques with intranasal administration. We produced three AAVs expressing with GFP or RFP using the CAG promoter (i.e., AAV9-RFP, AAV9-GFP, and AAV.CPP.16-GFP) and verified their infectivities in HEK293T cells in vitro (**Supplementary Fig. S3a,b**). Three NHPs were prepared after pre-screening for neutralizing antibodies against AAV9 capsid (Antibody titer <1:10, **Supplementary Fig. S3c**). Serving as a control, AAV9-RFP was co-injected in one NHP (#1) with AAV9-GFP and in two NHPs (#2 and #3) with AAV.CPP.16-GFP (4 × 10^12^ vg per AAV per animal). 4 weeks after AAV administration, all 3 NHPs were subjected for histological examination. No behavioral abnormality or weight loss was observed in any of the three animals. Comprehensive blood tests performed across the duration of the study revealed no major adverse effects associated with AAV administration (See **Supplementary Table S1-3**).

In NHP #1 treated with AAV9-GFP/AAV9-RFP co-administration, a small amount of RFP and GFP labeled cells were observed on sections sampled from the nasal cavity, trachea and lung (**Fig. 3a,c,e**). Cells co-labeled with RFP and GFP were not detected. For quantification, ImageJ software was used to measure the RFP and GFP fluorescence intensities using DAPI nuclear staining signal as benchmark. The GFP/RFP ratio was calculated and used for comparing the transduction efficiencies of AAVs. As expected, AAV9-GFP and AAV9-RFP had similar transduction efficiencies for all examined tissues with the GFP/RFP ratios close to 1 as seen in NHP#1.

**Fig. 3.**
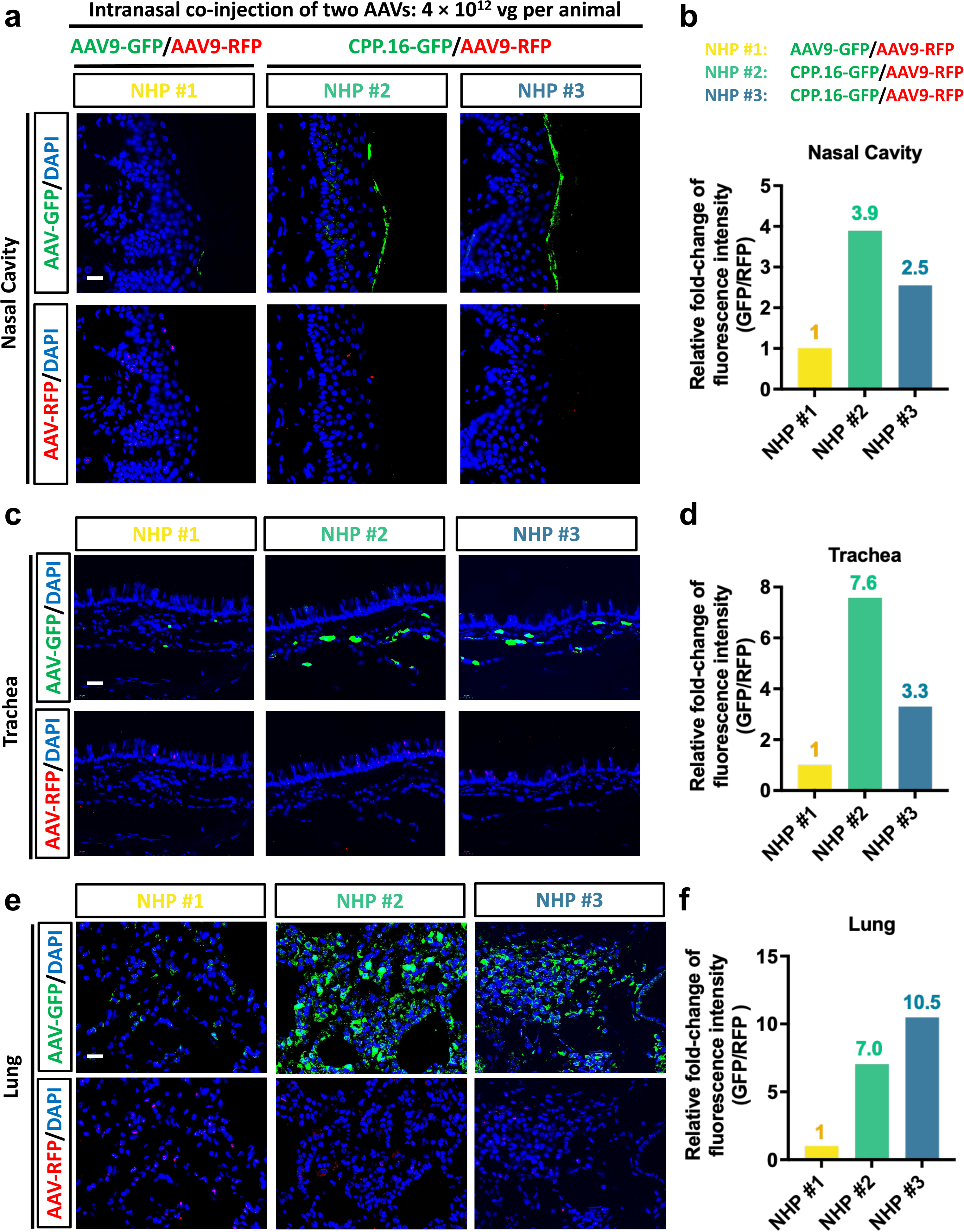
Intranasal AAV.CPP.16-mediated gene delivery in NHPs. **a, c, e,** Representative images showing AAV-mediated GFP and RFP co-expression in the nasal cavity (a), trachea (c) and lung (e) in three adult NHPs. DAPI nuclear staining was shown in blue. In NHP #1, AAV9-GFP and AAV9-RFP were pre-mixed and co-administered intranasally. In NHPs #2 and 3, CPP.16-GFP and AAV9-RFP were co-administered. AAV9-RFP serves as an internal control. 4 × 10^12^ vg of AAVs per AAV per animal were applied. Tissues were collected 28 days after AAV administration. Scale bar: 20 μm. **b, d, f,** Quantification of GFP fluorescence intensity measured as the ratio of GFP/RFP in the nasal cavity (b), trachea (d) and lung (f). Values were normalized to those in NHP#1 to show the fold changes of CPP.16-mediated GFP expression relative to AAV9.

In NHPs #2 and #3 treated with AAV.CPP.16-GFP/AAV9-RFP co-administration, substantially stronger GFP fluorescence was measured in the respiratory tract in comparison with the RFP signal, suggesting a better transduction efficiency by AAV.CPP.16 relative to AAV9. Change folds of AAV.CPP.16 vs. AAV9 ranged from 3.9 to 7.6 in NHP#2 and from 2.5 to 10.5 in NHP#3 in examined respiratory parts (**Fig. 3b,d,f**). As in the mouse study, we further examined the cell types transduced by the AAVs including MUC5A-positive goblet cells in the nasal cavity, CC10-positive club cells in the trachea, and RAGE-positive alveolar type I cells in the lung. While AAV9 resulted in little transduction of these three cell types, AAV.CPP.16 was able to transduce 23% and 14% of goblet cells, 11% and 18% of club cells, 33% and 27% of alveolar type I cells, in NHP#2 and NHP#3 respectively (**Supplementary Fig. S3d,e,f**). These results suggest the enhanced tropism of AAV.CPP.16 over AAV9 can translate from mice to NHPs.

Additional histology analysis in all three NHPs revealed no detectable AAV expression in the heart, kidney, spleen or brain tissues (**Supplementary Fig. S4,S5**). As shown in mice, we found AAV.CPP.16-GFP resulted in the transduction of some cells in the liver in both NHP#2 and NHP#3 (**Supplementary Fig. S4**).

### AAV.CPP.16-mediated CRISPR gene therapy against SARS-CoV-2

Having observed the enhanced gene delivery efficiency of AAV.CPP.16 in both proximal and distal airways and the lung, we wanted to explore its potential application in gene therapy for respiratory diseases. The continuing emergence of potentially more dangerous SARS-CoV-2 variants is concerning. The CRISPR strategy has the potential of generating a broad-spectrum therapy by targeting critical conserved genomic regions of all SARS-CoV-2 variants; it may be a valuable approach to counteracting rapid evolution and mutation of SARS-CoV-2 and provide a complementary therapy in addition to the current vaccination and antiviral drug treatments. However, the safe and efficient delivery of a CRISPR gene therapy to the respiratory tract has been a major technical barrier. Here, we designed a new “all-in-one”, broad-spectrum CRISPR gene therapy against SARS-CoV-2 and explored the application of AAV.CPP.16 for its in vivo delivery.

For targeting viral RNA in SARS-CoV-2, we chose to use CasRx, a RNA-guided CRISPR RNA endonuclease that belongs to the class 2 type VI-D CRISPR-Cas13d system. Derived from Ruminococcus flavefaciens XPD3002 (RfxCas13d, or CasRx)^19^, CasRx is small in size (967 amino acids), specific and strong in its catalytic activity; it employs short, customizable CRISPR-associated guide RNAs (gRNAs) for target recognization and engagement. To design effective gRNA sequences, we first performed genome mapping of sequenced SARS-CoV-2 strains and analyzed CasRx-targetable regions. We analyzed 2815 SARS-CoV-2 genomes downloaded from the public database NCBI SARS-CoV-2 Resources (http://www.ncbi.nlm.nih.gov/genbank/sars-cov-2-seqs/), and identified 7 conserved regions located within putative open reading frames, including regions 3C-like protease (3CL), Programmed ribosomal frameshifting (PRF), RNA-dependent RNA polymerase (Rdrp), Spike (S), Envelop (E), Membrane (M), and Nucleocapsid (N) (**Fig. 4a**). The conservation scores of these 7 regions arranged from 0.973 to 0.999 out of 1 (**Fig. 4b, Supplementary Table S4**), suggesting vital roles of these genes and thus potential value serving as targets. We then performed 23-nucleotide (nt) candidate gRNAs prediction analysis for all 7 conserved genes based on published Cas13 gRNA design principles^20^ and generated standardized guide scores for all candidate gRNAs (**Supplementary Fig. S6**). gRNA sequences that belonged to the 4^th^ quartile were predicted to have good targeting efficiency. We decided to focus on the Rdrp as the CasRx targeting gene because: 1, sufficient output of good gRNA candidates was obtained with 460 candidates located at the 4^th^ quartile out of 2,614 sequences in total; 2, Rdrp is essential for the replication of the SARS-CoV-2 viral genome and transcription of its genes^21,22^, and its value as a therapeutic target has been demonstrated by several clinically proven small molecule antiviral drugs^23–26^; 3, Rdrp is believed to be under less evolution pressure, as compared with the Spike gene, so the risk of mutational escape would be lower.

**Fig. 4.**
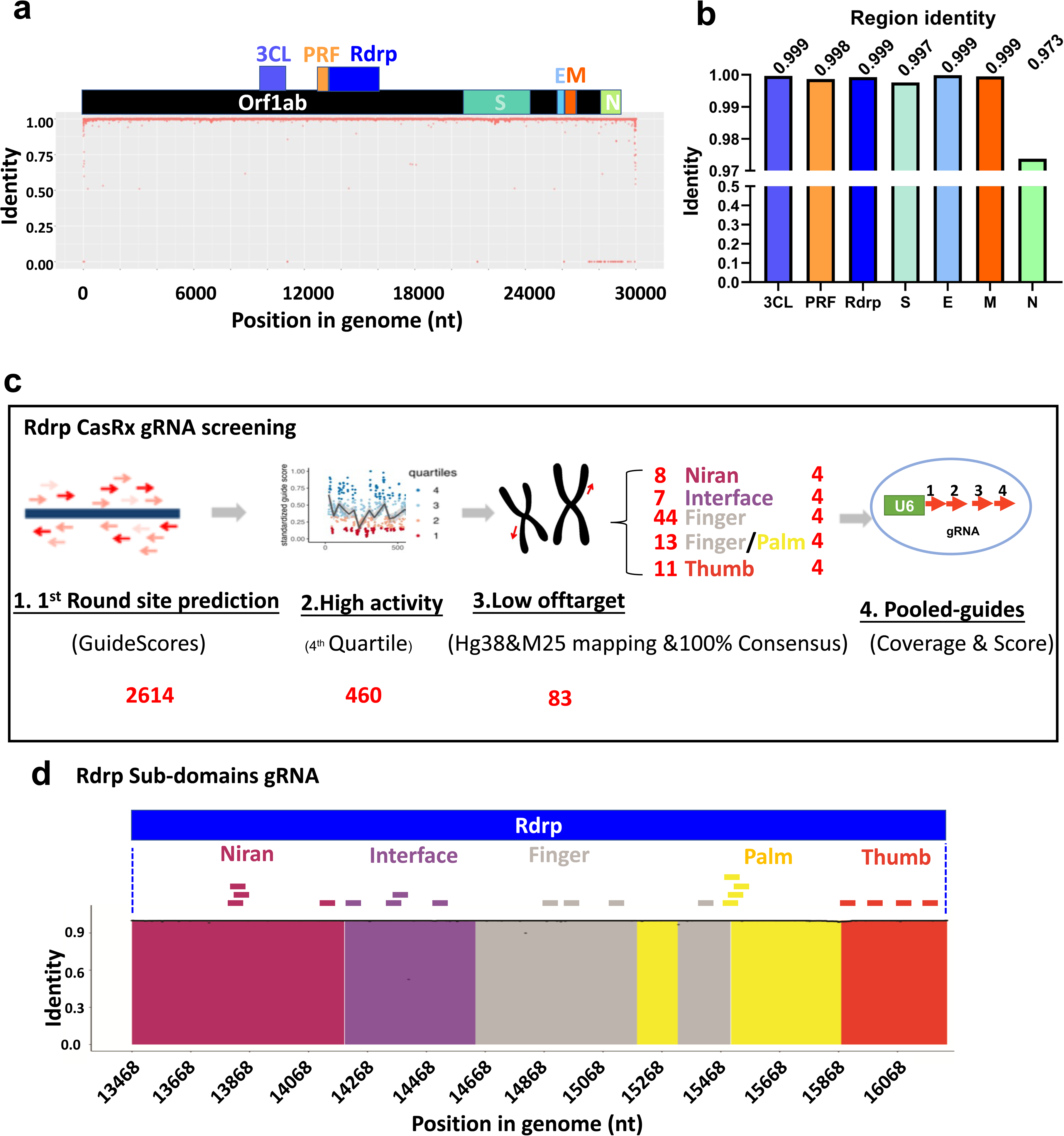
Identification of CasRx target sites in SARS-CoV-2 genomes. **a**, Alignment of 2815 patient-derived SARS-CoV-2 sequences downloaded from public database NCBI SARS-CoV2 Resources. 7 regions are highlighted including 3C-like protease (3CL), Programmed ribosomal frameshifting (PRF), RNA-dependent RNA polymerase (Rdrp), Spike (S), Envelop (E), Membrane (M) and Nucleocapsid (N). **b**, Conservation of selected genes across SARS-CoV-2 genomes. **c**, Workflow for selecting CasRx gRNAs that target the Rdrp gene. **d,** schematic diagram showing targeting sites in the Rdrp gene for all selected gRNAs. The Rdrp gene encodes for the 932-aa Rdrp protein that contains four conserved regions including Niran, Interface, Finger, Finger/Palm and Thumb.

We performed off-target filtration on the 460 gRNAs with high predicted Rdrp-targeting scores, excluded 377 of them with off-targets containing 2 or fewer mismatches in the human and mouse transcriptomes, and retained 83 with little predicted off-targeting. Four conserved regions are previously described in the Rdrp protein including Niran, Interface, Finger, Finger/Palm and Thumb^21,23,27^. We found 8 of the 83 remaining gRNAs target the Niran region, 7 target Interface, 44 target Finger, 13 target Finger/Palm,11 target Thumb (**Fig. 4c**). We selected 4 gRNAs with relatively wide coverage for each conserved region of Rdrp and constructed a pooled 4x gRNA sequence that contains all 4 gRNAs in tandem. 5 pooled gRNA sequences were generated in total with each targeting one of the five conserved regions in Rdrp (**Fig. 4d and Supplementary Table S5**).

Next, we designed and constructed a dual-promoter, all-in-one AAV-CasRx-gRNA system that could express both CasRx and the 4x gRNAs in one vector (**Fig. 5a and Supplementary Table S6**). CasRx with two 7-amino acid SV40 nuclear localization signal (NLS) sequences and one 9-amino acid HA tag was expressed under the regulation of the small EFS promoter (212-bp or base pairs) and a 2x sNRP1(the soluble neuropilin-1) polyadenylation sequence (34 bp)^28^. The U6 promoter was used to drive the expression of the 4x gRNAs and the 30-bp Cas13d direct repeats (DRs) that are essential for gRNA expression^19^. The length of the DNA sequence between the two inverted terminal repeats (ITRs) was approximately 3.8k bp, which is well within the 4.7k bp packaging capacity for AAV. With five different 4x gRNA sequences targeting the Rdrp, we generated five AAV plasmid vectors including pAAV-CasRx-gRNA.Niran, pAAV-CasRx-gRNA.Interface, pAAV-CasRx-gRNA.Finger, pAAV-CasRx-gRNA.Palm, and pAAV-CasRx-gRNA.Thumb. In addition, we generated two control plasmids by replacing the 4x gRNA with a scramble sequence (pAAV-CasRx-scramble) and a previously published gRNA sequence targeting the Rdrp (pAAV-CasRx-gRNA.Abbott)^22^. To test the Rdrp inhibition efficiency of our plasmid vectors, we co-transfected each of the pAAV-CasRx plasmid vectors into HEK-293T cells with a Rdrp reporter plasmid and measured Rdrp transcript abundance RNA 48 hours later by quantitative real-time PCR (**Fig. 5b**). Compared with Rdrp transfection alone, co-transfection of the scramble plasmid resulted in no reduction of Rdrp transcription. Similar to the effect of the “Abbott” plasmid which deceased Rdrp by 46%, all our Rdrp-targeting plasmid vectors achieved Rdrp inhibition by a magnitude ranging from 44% to 58% (**Fig. 5c**).

**Fig. 5.**
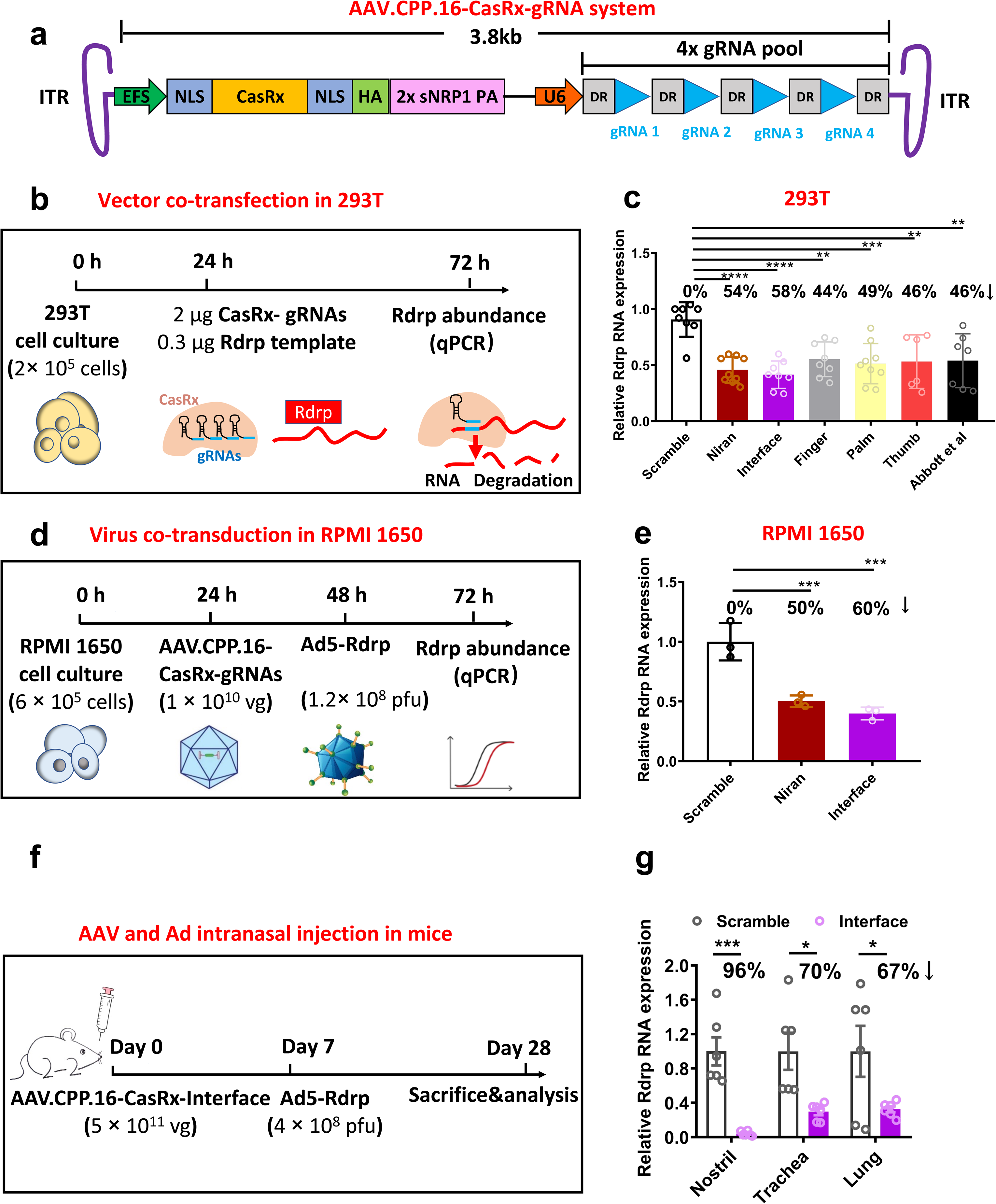
CasRx-mediated knock-down of Rdrp transcription using an all-in-one AAV.CPP.16 vector. **a**, Schematic diagram showing an all-in-one AAV vector expressing both CasRx and four gRNAs. **b**, Workflow for testing Rdrp RNA knock-down in 293T cells by co-transfection with CasRx-gRNA and Rdrp plasmids. **c**, Quantification of Rdrp knock-down after co-transfection in 293T. Abbott et al: gRNA from a published study^22^. n = 6-9 per group, mean ± s.e.m., one-way ANOVA with Tukey’s post-test. **d**, Workflow for testing Rdrp RNA knock-down in RPMI 1650 cells by co-transduction with AAV.CPP.16-CasRx-gRNA and Ad5-Rdrp. **e**, Quantification for knock-down of Ad5-expressed Rdrp transcript by AAV.CPP.16-CasRx-gRNA in RPMI 1650 cells. n = 3 per group, mean ± s.e.m., one-way ANOVA with Tukey’s post-test. **f**, Timeline of studying intranasal AAV.CPP.16-CasRx-Interface gRNA and Ad5-Rdrp virus challenge in mice. **g**, Quantification of Rdrp knock down in the nostril, trachea and lung of mice. n = 6 each group, mean ± s.e.m., two-way ANOVA with Sidak’s post-test. *p < 0.05, **p < 0.01, ***p < 0.001, ****p < 0.0001.

Next, we tested AAV.CPP.16-mediated Rdrp inhibition as a prophylactic treatment strategy against SARS-CoV-2 in a pseudovirus model. Recombinant Ad5 viruses carrying the viral Rdrp gene (1.2×10^8^ pfu/well) were used to artificially express the Rdrp gene in the human nasal epithelial cells RPMI-1650 (**Fig. 5d**). Two most efficient AAV plasmid vectors (i.e., “Niran” and “Interface” with inhibition efficiencies of 54% and 58% respectively), along with the scramble, were selected and packaged into AAVs with the AAV.CPP.16 capsid. Indeed, treatment of AAV.CPP.16-CasRx-gRNA.Niran or AAV.CPP.16-CasRx-gRNA.Interface (5×10^11^ vg/ well) prevented Ad5-mediated Rdrp transcription by 50% and 60% respectively (**Fig. 5e**). For further testing in mice, intranasal administration of AAV.CPP.16-CasRx-gRNA.Interface or AAV.CPP.16-CasRx-scramble (5×10^11^ vg/ mice) was performed in adut C57BL/6 mice, followed by a second intranasal administration of Ad5-Rdrp (4×10^8^ pfu/mice) 7 days later (**Fig. 5f**). Animals were sacrificed after another 3 weeks and tissues collected for RNA analysis. Ad5-Rdrp alone resulted in remarkable over-expression (50~300 fold) in mouse nostril, trachea and lung 4 weeks after intranasal injection (**Supplementary Fig. S7**). Compared with the scramble AAV, AAV.CPP.16-CasRx-gRNA.Interface resulted in significantly reduced Rdrp transcription in all examined regions of the respiratory tract with near complete blockade of Rdrp transcription in the nostril (**Fig. 5g**).

### Broad-spectrum protection for SARS-CoV-2 variants

SARS-CoV-2 is constantly evolving and accumulating mutations in its genetic code. Genomic epidemiology of SARS-CoV-2 has changed significantly since the pandemic outbreak in 2019 (**Fig. 6a**). As the Rdrp-targeting interface-gRNAs were designed based on genomic sequences of early SARS-CoV-2 lineages (Wuhan-Hu-1/ 2019), we did a comprehensive mutation analysis on the Rdrp gene across all sequenced SARS-CoV-2 lineages and examined the theoretical compatibility of the “interface” gRNAs. Based on the World Health Organization (WHO) classification, 10 major SARS-CoV-2 variant categories have been described including Alpha, Beta, Gamma, Delta, Eta, Iota, Kappa, Lambda, Mu, and Omicron. Each WHO category may include several lineages as designated using the Phylogenetic Assignment of Named Glocal Outbreak (PANGO) nomenclature^29^. We thus analyzed all 10 WHO categories and 12 PANGO lineages using the online database https://outbreak.info/. Compared with the origin strain Wuhan-Hu-1/ 2019^30^, only three mutations P314L, P314F and G662S were found in the Rdrp gene (**Fig. 6b and Supplementary Table S7**). This is in stark contrast to hundreds of mutations observed in the Spike protein^31^. We next aligned the 4 interface-gRNAs with all examined genomes and found a perfect match between any of the gRNAs and its target sequences (**Fig. 6b, c and Supplementary Table S8**). These results suggested a potentially broad-spectrum efficiency of our interface-gRNA-guided CRISPR gene therapy for both circulating and emergent SARS-CoV-2 variants.

**Fig. 6.**
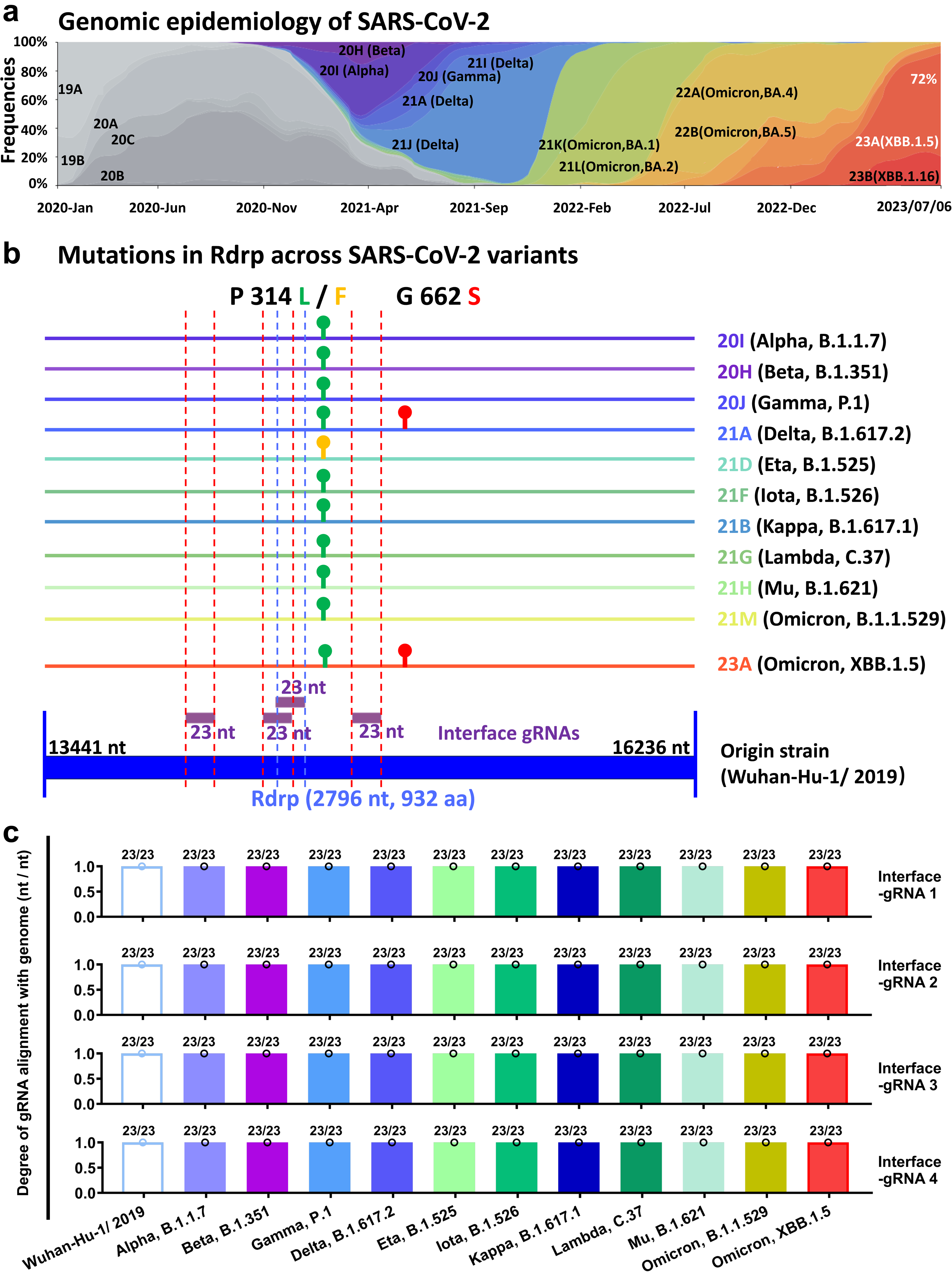
Projected targeting of Interface-gRNAs across all known SARS-Cov-2 variant categories. **a**, Globally genomic epidemiology of SARS-CoV-2 since COVID-19 outbreak. **b**, Mutations in the Rdrp gene across representative SARS-Cov-2 variants as compared with the origin strain Wuhan-Hu-1/2019. All mutations are located outside the targeting regions of the four 23 nt Interface-gRNAs. **c**, Quantification of four Rdrp-Interface gRNAs alignment with 11 variants genome (nt / nt). No mismatch was observed between any gRNA and viral genome.

## Discussion

### AAV gene delivery for the respiratory system

It has proven to be more challenging than originally anticipated to develop gene therapy for the treatment of respiratory diseases such as cystic fibrosis, or against airborne viral infection. Efficient delivery of therapeutic genes to disease-relevant cell types, mainly the airway epithelial cells and lung parenchyma cells, remains a major technical hurdle. With fast-accumulating clinical experience including several regulatorily approved drugs, AAV has its advantages as a widely used gene delivery modality such as a relatively good safety record and versatility in targeting a variety of tissue and cell types. To develop efficacious gene therapy for respiratory conditions, most likely, by intranasal administration, a primary goal is to achieve an adequate percentage of transduction in both proximal and distal airway cells. AAV6 and AAV9 are among the most efficient wild-type AAV serotypes targeting the respiratory tract and the lung based on studies in both small and large animal mdoels^32^. However, overall transduction efficiency by these two AAVs is still sub-optimal especially for large animals such as the NHPs, limiting their clinical translation^4^. AAV.CPP.16 is an engineered capsid derived from AAV9 by inserting a 6-mer short peptide into its VP capsid proteins^6^. Although AAV.CPP.16 was designed as a BBB-penetrant capsid for the CNS, it was observed that it also transduced more cells in the lung than AAV9 after systemic administarion. This led us to further assess it for potential application of intranasal gene delivery targeting the respiratory airways. Indeed, AAV.CPP.16 outperforms AAV6 and AAV9 in transducing the cultured human nasal epithelial cells and all examined mouse airway and lung cells. The observed superiority of AAV.CPP.16 over control AAV (i.e., AAV9) is particularly notable in our NHP studies. Although more studies are warranted, for example, on more characterization of AAV.CPP.16 in a larger NHP study, our results suggest AAV.CPP.16 as an improved capsid for targeting the airway and the lung with potential value of translation.

To address the issue of high turnover of airway epithelial cells and thus achieve sustained therapeutic effect of AAV gene therapy, two AAV-mediated approaches have been successfully tested in animal models: CRISPR-based genome editing that makes inheritable genetic changes^17^ and repeated administration using AAV9 as capsid^3^. Given its enhanced targeting efficiency, it would be interesting to test AAV.CPP.16 for delivering CRISPR genome editing machinery for the treatment of hereditary respiratory diseases such as cystic fibrosis. Although AAV.CPP.16 is derived from AAV9, it would also be of importance to test (and confirm) whether repeated dosing of AAV.CPP.16 is feasible.

### CRISPR-based gene therapy against SARS-CoV-2

CRISPR-Cas13 is a versatile tool for RNA knock-down and binding^33^, and has been used for RNA virus detection and inhibition^34,35^. Thus, CRISPR-Cas13 could be a promising antiviral strategy for broadly targeting SARS-CoV-2 variants. Indeed, following the outbreak of COVID-19 in 2020, an editorial perspective proposed the concept of “virus against virus” and suggested a roadmap of using an AAV-based CRISPR/Cas13d system to combat SARS-CoV-2, although no experimental test was performed^36^. Subsequently, several studies explored CRISPR-Cas13 as an antiviral strategy against SARS-CoV-2, influenza and other human-injecting RNA viruses and tested the feasibility of such strategy using *in vitro* assays^22,37,38^. Our study has extended previous findings by using CasRx, a member of the Cas13 family to target the well-conserved Rdrp protein in SARS-CoV-2. We provide both *in vitro* and *in vivo* efficacy evidence supporting further development of CRISPR-Cas13 against SARS-CoV-2 using a clinically relevant AAV vector AAV.CPP.16. Our leading candidate therapy (i.e., AAV.CPP.16-CasRx-gRNA.Interface) knocks down Ad5-mediated Rdrp transcription by more than 60% in human nasal epithelial cells and mouse respiratory tract tissues. Intranasal pre-treatment of such gene therapy in mice almost totally prevents Ad5-mediated Rdrp transcription in the nostril, a key anatomical location for initial viral infection and application when SARS-CoV-2 targets human.

One limitation of our study is that it remains unknown whether our AAV.CPP.16-CRISPR therapy has any antiviral effect against real SARS-CoV-2 viruses. Follow-up research will focus on testing the candidate therapy against SARS-CoV-2 challenge in cultured cells and in a human-ACE2 transgenic mouse line. Despite success of vaccination and other strategies targeting SARS-CoV-2, the risk of “breakthrough” SARS-CoV-2 mutants exists and treatment options for immune-compromised patients are still limited. The high efficiency of AAV.CPP.16 in targeting respiratory tract tissues in primates, robust and broad spectrum of our CRISPR gRNA array, and versatility of the CRISPR system in general all warrant further development of our gene therapy as a supplementary strategy against current and emerging SARS-CoV-2 variants.

In summary, we describe AAV.CPP.16 as an intranasal gene delivery vector with high tropism for respiratory tract tissues in both mice and non-human primates. We also report an AAV.CPP.16-based CRISPR-Cas13 gene therapy with potentially broad antiviral effects against SARS-CoV-2 variants.

## Methods

### Cell culture

All cell lines were incubated at 37 °C and at 5% CO_2_. Human embryonic kidney (HEK293T) cells were cultured in 10% fetal bovine serum (Thermo Fisher Scientific) in DMEM (Cat# 11995073, Gibco). Human nasal epithelial cell RPMI 1650 (CCL-30, ATCC) were cultured in 10% FBS in EMEM (30-2003, ATCC).

### Animals

Mouse experiments were approved by the Institutional Animal Care and Use Committee (IACUC) at Brigham and Women’s hospital, Ninth People’s Hospital, Shanghai JiaoTong University School of Medicine, and Soochow University. For AAV distribution study, both male and female adult mice aged over 8 weeks were used. Mouses strains C57BL/6J were purchased from Jackson Laboratory.

Non-human primate (NHP) study was performed by the Association for Assessment and Accreditation of Laboratory Animal Care (AAALAC)-accredited contract research organization Kunming Primate Research Center (Kunming Institute of Zoology, Chinese Academy of Sciences, Kunming, China) with its IACUC approval. All NHP protocols were further reviewed and approved by Kunming Medical University. NHPs (*Cynomolgus macaques*) were pre-screened for health status before study enrollment and housed in individual cages on a 12-h light/dark schedule in well ventilated rooms (temperatures:18-29 °C, humidity: 40-70%). Standard care was provided to all NHPs in accordance with AAALAC and local research animal guidelines. All NHPs were screened against pre-existing neutralizing antibody of AAV9.

### AAV production

Recombinant AAVs used for experiments other than in NHP were packaged in-house as previously reported^6^. Briefly, the HEK293T cells were co-transfected with three-plasmids, RC plasmid containing AAV rep and cap genes, pHelper plasmid (240071-12, Agilent Technologies) and pAAV for in a 1:1:1 molar ratio using polyethylenimine (PEI, Cat# 23966, Polysciences). The AAV production media were collected 72 h and 120 h post transfection and concentrated using a PEG-precipitation method with 8% PEG-8000 (wt/vol; Cat# V3011, Promega). The harvested cell pellets were re-suspended and lysed through sonication. Vectors were purified by iodixanol gradient (15%, 25%, 40% and 60%) with ultracentrifugation at 48,000 x *g* for 1 h at 18°C. rAAVs were then buffer-exchanged and concentrated using Millipore Amicon filter unit (UFC910008, 100K MWCO) with 1x DPBS containing 0.001% Pluronic F-68 (Gibco). AAV titer was quantified by measuring DNase-resistant genome copies using “Free-ITR” qPCR method with specific primers detecting WPRE element in AAV vectors. All titers were calculated using PVUII (NEB) digested linearized AAV plasmid as a standard and normalized to a rAAV-2 reference standard (Cat# VR-1616, ATCC). AAVs used in NHP studies were produced and tittered by PackGene Biotech.

### In vitro AAV transduction assay

For RPMI 1650 cells, 4 × 10^9^ vg of AAV-CAG-GFP was applied at MOI of 1 × 10^5^ in 24-well plates. For 293 cells, 1 × 10^10^ vg of AAVs were applied at MOI of 5 × 10^4^. 72 hours after AAV infection, cells were fixed with 4% PFA for 15 min and followed by DAPI staining for 30 min. Intensity of native fluorescence was measured using ImageJ and normalized to DAPI signal.

### Intranasal administration in mice

Adult mice older than 8 weeks of age were anaesthetized with 2.5% isoflurane and placed in a prone position. A blunt needle attached to a 10 µL hamliton syringe (Cat # Model 701N, Hamilton) was inserted into the back of one nostril. 20 µL of AAV vectors diluted in sterile saline were slowly infused while ensuring no airway blockade. A second infusion of 20 µL of AAV solution was performed in the other nostrila to add up to 40 µL of AAV vectors per animal. Similar procedure was performed for intranasal administration of Ad5. For tissue harvest, transcardial perfusion was performed with PBS before tissues were collected for viral genome quantification, or with PBS and then followed tissue fixation using 4% paraformaldehyde (PFA) for histology use.

### Intranasal administration in NHPs

NHPs (*Cynomolgus macaques*) were sedated with ketamine (10 mg/kg). AAV vectors diluted in 1x PBS were slowly injected into the nostril using a 1 mL pipette. The total injection volume per animal was 1 mL with 0.5 ml for per nostril. Blood samples were taken 1 day before AAV injection (day 0), 3 days and 21 days after AAV injection for metabolic panel test and complete blood count. Animals were monitored closely, checked at least twice a day for any signs of abnormality. No immunomodulatory drug was administered. 3 weeks after AAV injection, animals were anesthetized with phenobarbital sodium (30 mg/kg) and subjected to transcardial perfusion with PBS, followed by 4% PFA. Tissues were then harvested and processed for paraffin embedding and sectioning.

### Neutralizing antibody titration

Neutralizing antibody titers in NHPs were determined using a cellular luciferase report assay. Venous blood samples were collected into serum-separating vacutainer tubes (BD). Serum supernatant was isolated by centrifuging at 1,000x g for 10 mins and transferred to a clean tube. HEK293T cells were seeded at 2 × 10^4^ per each well in 96-well plates (Corning). 24 hours later, serial-diluted serum samples were prepared and mixed with AAV9 expressing luciferase (10^4^ vg/cell). The mixtures were then incubated at 37 °C for 1 hour prior to adding to the prepared HEK293T cells. 24 hours later, the cells were prepared for the Bright-Glo Luciferase Assay according to the manufacturer’s protocol (Promega). Luciferase activity was detected by using a luminescence microplate reader (Cytation 5, BioTek). Neutralizing antibody of selected NHPs was detected as 1:10 dilution at which 50% or less inhibition of the luciferase signal was measured.

### Histology and immuno-staining

For mice, PFA-fixed tissues were dehydrated in 30% sucrose and embedded in OCT for frozen sectioning. Typically, 30 µm thick sections for tissues were cut for standard immunohistochemistry with antibody staining.

Processing of NHP tissues was carried out by Servicebio Inc (Wuhan, China). Briefly, tissues were fixed in a 4% paraformaldehyde (PFA) solution for over 24 hours. Sections with a thickness of 20 µm were prepared, deparaffinized using xylene, and rehydrated through a series of ethanol gradients. Antigen retrieval was conducted by employing a citric acid (pH 6.0) antigen repair solution at a sub-boiling temperature. Following this, the sections were immersed in a 3% hydrogen peroxide solution to inhibit endogenous peroxidase activity, and subsequently incubated in a 3% bovine serum albumin solution to prevent nonspecific binding. For multiple immunofluorescent co-labelling, tyramide signal amplification (TSA) protocol was used. First, sections were incubated with the anti-RFP antibody overnight, and followed by incubation with a horseradish peroxidase (HRP)-conjugated secondary antibody (1:500, GB23303, Servicebio) for 50 mins at room temperature in dark condition. Then, slides were further treated with a CY3-TSA solution (G1223, Servicebio) for 10 mins in a dark condition. After that, both first primary and secondary antibodies were removed through the application of an EDTA antigen repair solution. To detect GFP signal on the same slides, a mouse anti-GFP antibody was applied, followed by applicaton of a Alexa Fluor 488-conjugated goat-anti-mouse secondary antibody (1:400, GB25301, Servicebio). To visualize additional antigens including cell type markers, slides were further incubated with relevant primary antibodies and then with corresponding HRP-conjugated secondary antibodies, before being treated with a CY5-TSA solution. Nucleus counterstaining was performed using DAPI. Sections were imaged using a slide scanner (PANNORAMIC, 3D HISTECH).

Primary antibodies included: chicken anti-GFP (1:1000, AB13970, Abcam), mouse anti-GFP (1:3000, GB12602, Servicebio); rabbit anti-RFP (1:1000, 600-401-379, Rockland), rabbit anti-α-Tubulin (1:400, 5335, Cell signaling technology), rabbit anti-α-Tubulin (1:200, D20G3, Cell signaling technology), mouse anti-MUC5A (1:400, MA5-12178, Invitrogen), mouse anti-CC10 (1:400, sc-390313, Santa Cruz), rabbit anti-CC10 (1:200, A16997, abclonal), rat anti-RAGE (1:400, MAB1179-SP, R&D), rabbit anti-RAGE (1:200, 16346-1-AP, Proteintech). Fluorescence images were taken using confocal microscopy with integrated automatic image stitching function (Zeiss LSM710), processed and analyzed using ImageJ.

### SARS-CoV-2 genome sequences collection and conservation mapping

SARS-CoV-2 genome sequences were downloaded from NCBI SARS-CoV-2 Resources (http://www.ncbi.nlm.nih.gov/genbank/sars-cov-2-seqs/) on May 22,2020. Full genomes were mapped to SARS-CoV-2 Wuhan-Hu-1 NCBI reference genome (NC_045512.2, Genbank ID: MN908947.3, 29903 bp linear RNA) by MAFFT. The conservation identity was calculated as the percentage of sequences matching the consensus. The identity was determined by employing Geneious Prime (https://www.geneious.com) and subsequently visualized using R.

### CasRx gRNA design

To design gRNAs for SARS-CoV-2, the 99% consensus sequences of PRF, Rdrp, 3CL protease, S, E, M, and N, were imported into a CasRx design software (available at https://gitlab.com/sanjanalab/cas13) to predict potential guide sites and guide activity. 23-nucleotide (nt) sequences with perfect identity among the SARS-CoV-2 genomes were typically selected as candidates. Off-targets were scored against the human transcriptome (Hg38) and mouse transcriptome (M25) using Bowtie 1.2.3, to remove gRNAs that were mapped to the human and mouse transcriptome with less than 2 mismatches. 4 selected gRNAs were built into a gRNA pool for each targeting. Scramble and positive control sequence were referred to a previous study^22^.

### All-in-one CasRx-gRNA AAV vector

Plasmids pab202 and pab633 (gifts from Feng Zhang lab) and pAAV-CMV-dSa-VPR mini.-2X sNRP1 (a gift from George Church; Addgene plasmid # 99688) were used. Plasmid pab202, an AAV vector containing the EFS promoter, was used as AAV backbone. DNA sequences for NLS-CasRx-NLS-HA and 2x sNRP1 sequence (34 bp) were amplified from pab633 and pAAV-CMV-dSa-VPR mini.-2X sNRP1, respectively. The U6-DR-gRNA spacer fragment was synthesized by GenScript. Standard subcloning was performed to generate an “all-in-one” AAV vector (pAAV-EFS-CasRx-U6-4x gRNA).

### Rdrp reporter plasmid and Ad-Rdrp production

To express the Rdrp in HEK293T cells, the 2796 bp Rdrp genome sequence (NCBI nucleotide ID:NC_045512.2, 13441-16236) was synthesized and inserted into the pcDNA vector with a CMV promoter by GenScript. To express the Rdrp in RPMI 1650 cell and in mice, Rdrp was cloned into an Ad5-CMV vector (pADM-FH, Vigene Biosciences) and recombinant adenoviruses (Ad5-Rdrp) were packaged by Vigene Biosciences.

### Co-transfection in HEK-293T

2 × 10^5^ cells/well HEK-293T were plated in 24-well plates. On the following day, plamid pAAV-EFS-CasRx-U6-4x gRNA, Rdrp reporter plamid and PEI were mixed in a ratio of 2:0.3:6 in Opti-MEM. 48h after transfection, HEK-293T cells were harvested for RNA isolation.

### Co-transduction of AAV and Ad5 virus in RPIM-1650 cells

On day 0, 6 × 10^5^ cells/well RPMI-1650 cells were plated in 24-well plates. On day 1, RPMI-1650 cells were infected with 1 × 10^10^ vg AAV-CasRx-gRNA. On day 2, Ad5-Rdrp was added with a MOI of 200 (1.2× 10^8^ pfu). On day 3, RPMI-1650 cells were harvested for RNA isolation.

### Quantitative real-time PCR (qRT-PCR)

Quantitative real-time PCR was performed to quantify RNA abundance. For each sample, total RNA was isolated by using Trizol (Cat# 15596026, Invitrogen), followed by cDNA synthesis using the cDNA Synthesis Kit (Cat# 6110A, Takara). All qRT-PCR primers were ordered from Eton Bioscience. Quantitative real-time PCR was performed using the SYBR Green Mix (Cat# A25742, Applied Biosystems) in StepOne Plus system (Applied Biosystems) following manufacturers’ instructions. Ct values were used to quantify RNA abundance. The relative abundance of the Rdrp was normalized to a GAPDH internal control.

### Statistical analysis

All statistical analyses were performed using GraphPad Prism software. Two-sided Student’s t-test, one-way ANOVA with Tukey’s post-test, and two-way ANOVA with Sidak’s post-hoc tests were used for data comparison. p-values less than 0.05 were considered statistically significant.

## Acknowledgments

We thank Feng Zhang for sharing AAV and CasRx plasmids. This study is partly supported by a Brigham and Women’s Hospital sundry fund (FB).

## Author contributions

Z.Y. and Y.Z performed cell and mouse studies. Z.Y. and V.M. performed bioinformatic studies. X.C.and J.P. performed NHP study. X.F. contributed materials. F.B designed and superived studies. Z.Y., Y.Z. and F.B. analyzed data and wrote the manuscript.

