## Supplementary Figures for "Tropism of AAV.CPP.16 in the respiratory tract and its application for a CRISPR-based gene therapy against SARS-CoV-2"

### Supplementary information

#### Supplementary Figures

**Fig.S1.** related to Fig1 **| AAV transduction in brain, retina and multiple peripheral organs after intranasal administration.**

**Fig.S2.** related to Fig2 **| AAV-mediated transduction of mouse epithelial ciliated cells after intranasal administration.**

**Fig.S3.**related to Fig.3 **| AAV vector characterization, serum screening and transduction of cell type in NHP experiments.**

**Fig.S4.**related to Fig.3 **| AAV transduction in the heart, kidney, spleen and liver of the NHPs.**

**Fig.S5.**related to Fig.3 **| No AAV.CPP. 16 transduction in the NHP brain after intranasal injection.**

**Fig.S6.**related to Fig.4 **| Prediction of guide RNAs for conserved SARS-CoV-2 genes.**

**Fig.S7.**related to Fig.5 **| Expression of Rdrp in mice by intranasal administration of Ad5-Rdrp.**

#### Supplementary Tables

The Supplementary tables are provided as Additional Supplementary Information in the form of a single .xlsx spreadsheet in a separate file.

**Table.S1**, related to Fig.3 **| List of NHPs, AAVs administered and blood test results.**

**Table.S2**, related to Fig.3 **| Blood routine in all NHPs**.

**Table.S3.** related to Fig.3 **| Blood metaboic panel for all NHPs.**

**Table.S4.** related to Fig.4 **| SARS-CoV-2 RNA genome mapping.**

**Table.S5.** related to Fig.4 **| Selected guide RNAs targeting Rdrp.**

**Table.S6.** related to Fig.5 **| DNA sequences for AAV-CasRx-Rdrp Interface vector.**

**Table.S7.** related to Fig.6 **| Characteristic mutations in selected SARS-CoV-2 variants.**

**Table.S8.** related to Fig.6 **| Matching of Rdrp-Interface gRNAs in genomes of multiple SARS-CoV-2 lineages.**

**Fig. S1. related to Fig.1**

**a**

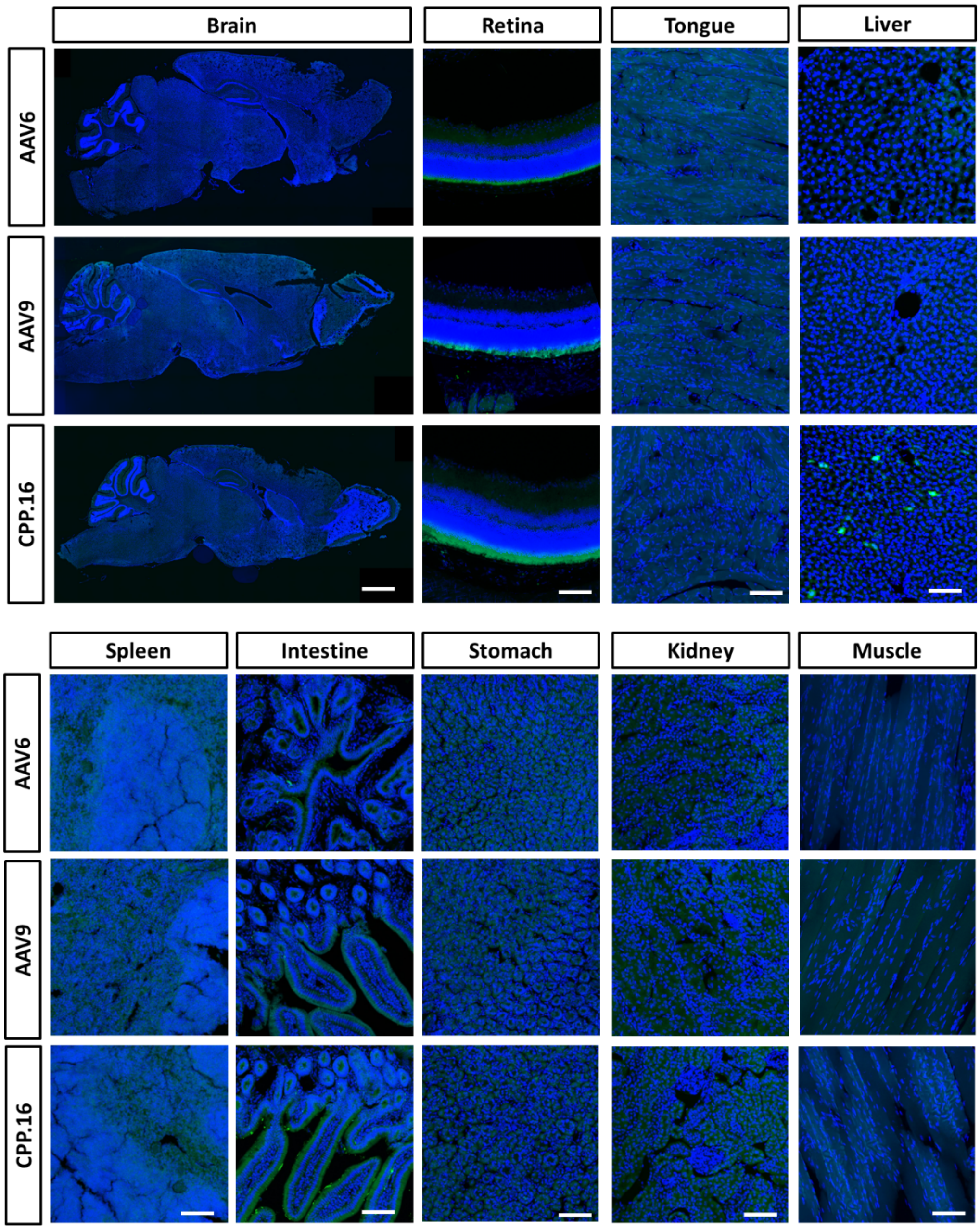


**Fig.S1.** related to Fig1 **| AAV transduction in brain, retina and multiple peripheral organs after intranasal administration.**

C57BL/6J mice received intranasal administration of 1 × $10^{11}$ vg of AAV6-CAG-GFP, AAV9-CAG-GFP, or AAV.CPP.16-CAG-GFP and were sacrificed 21 days later for harvesting tissues. Immunostaining with antibody against GFP was used to visualize transduced cells. No GFP-labeled cells were observed in the brain, retina, tongue, spleen, intestine, stomach, kidney or muscle. Some GFP-labeled cells were detected in the liver of AAV.CPP.16-treated mice. Scale bar: 200 μm for brain, 100 μm for other organs.

#### Fig.S2. related to Fig.2

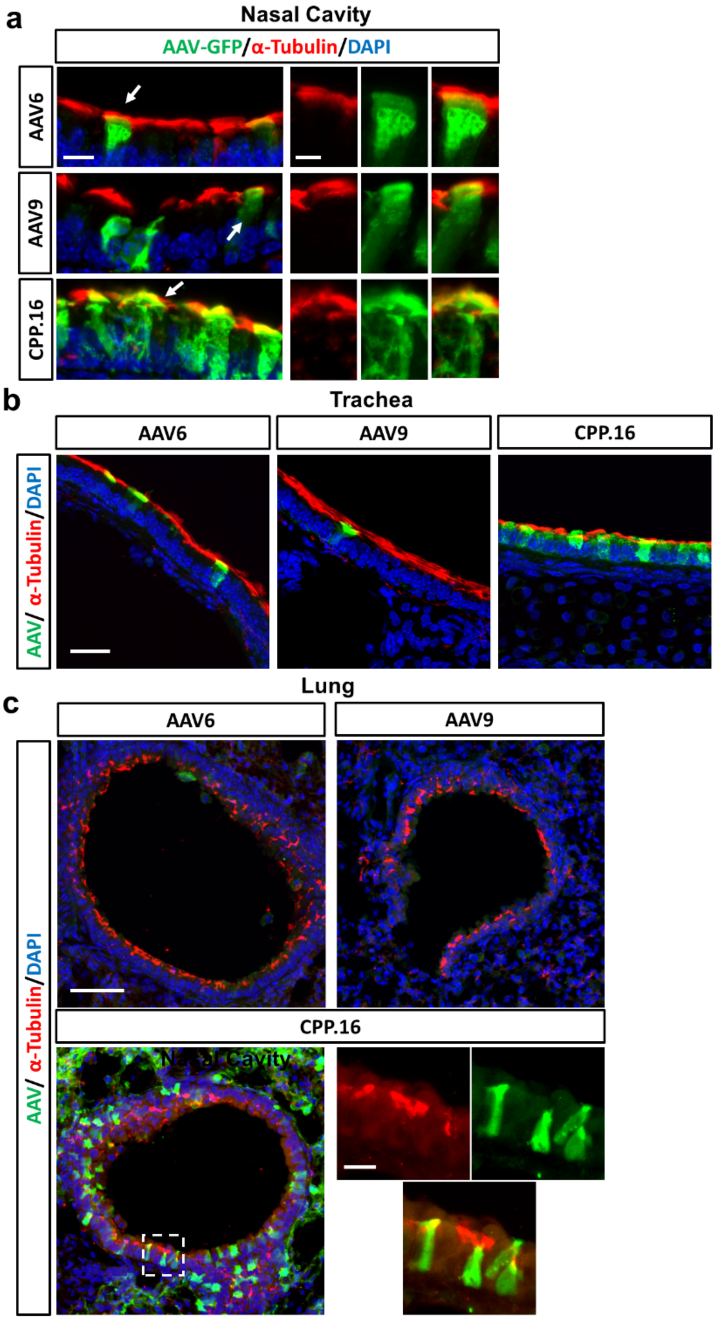


**Fig.S2.** related to Fig2 **| AAV-mediated transduction of mouse epithelial ciliated cells after intranasal administration.**

Representative images showing GFP-labeled ciliated cells in the nasal cavity (**a**), trachea (**b**), and lung (**c**) 21 days after intranasal administration of 1 × $10^{11}$ vg of AAV6-CAG-GFP, AAV9-CAG-GFP, or AAV.CPP.16-CAG-GFP. Ciliated cells were stained using an antibody against α-Tubulin. Scale bar in a: 10 μm for low-magnification images and 5 μm for enlarged images. Scale bar in b: 25 μm. Scale bar in c: 50 μm for low-magnification images and 10 μm for enlarged images.

**Fig.S3. related to Fig.3**

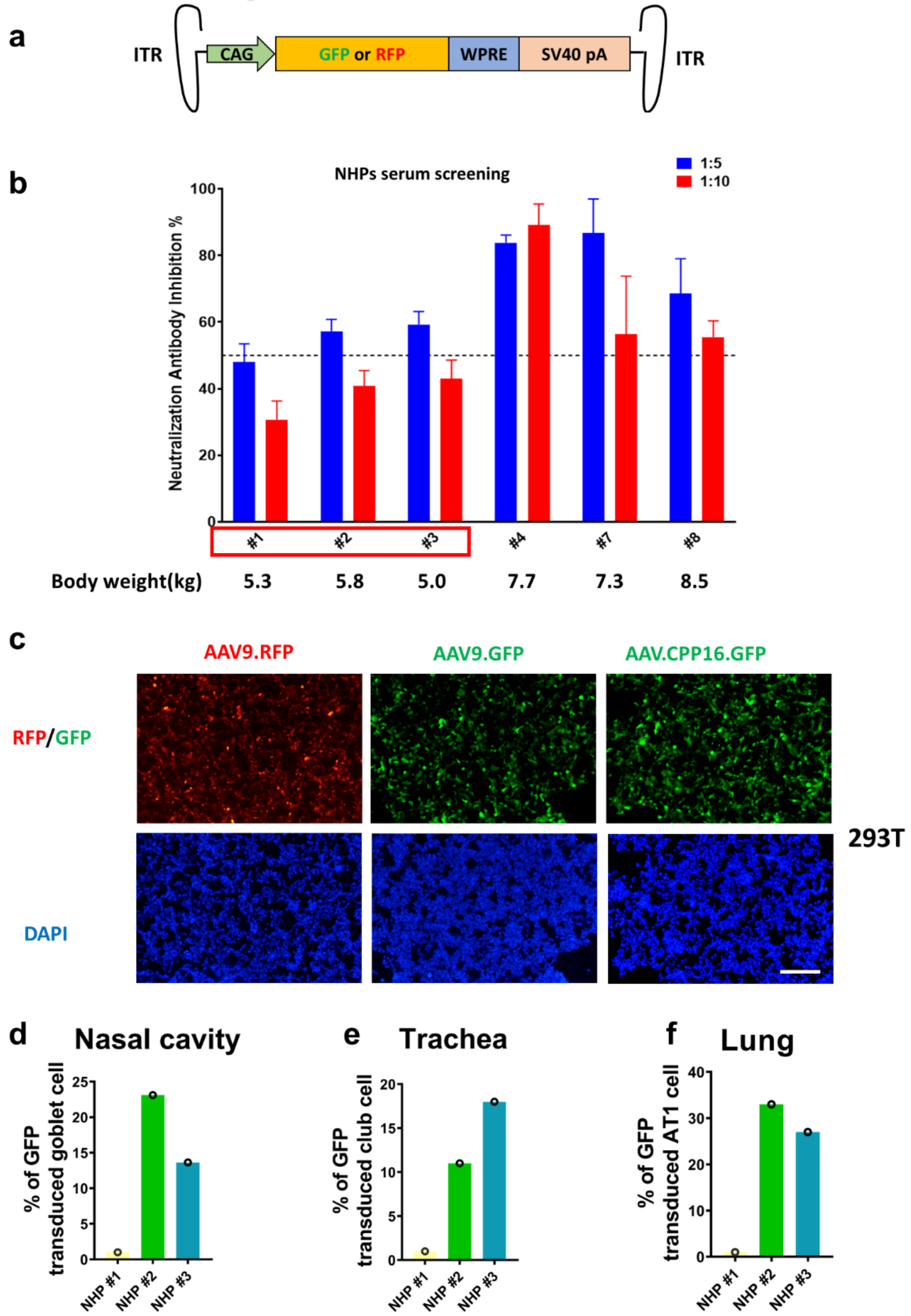


**Fig.S3.**related to Fig.3 **| AAV vector characterization, serum screening and transduction of cell type in NHP experiments.**

**a**, Diagram showing AAV vectors. **b**, *In vitro* transduction of AAV9-GFP, AAV9-RFP and AAV.CPP16-GFP in HEK293T cells. Scale bar 100 μm. **c**, Pre-screening of NHPs. NHPs were pre-screened against the pre-existing neutralizing antibodies of AAV9 (titer < 1:10).d-f, Percentage of AAV-GFP-transduced airway cells in NHP out of DAPI-labeled total cells. AT1, alveolar type 1.

#### Fig.S4. related to Fig.3

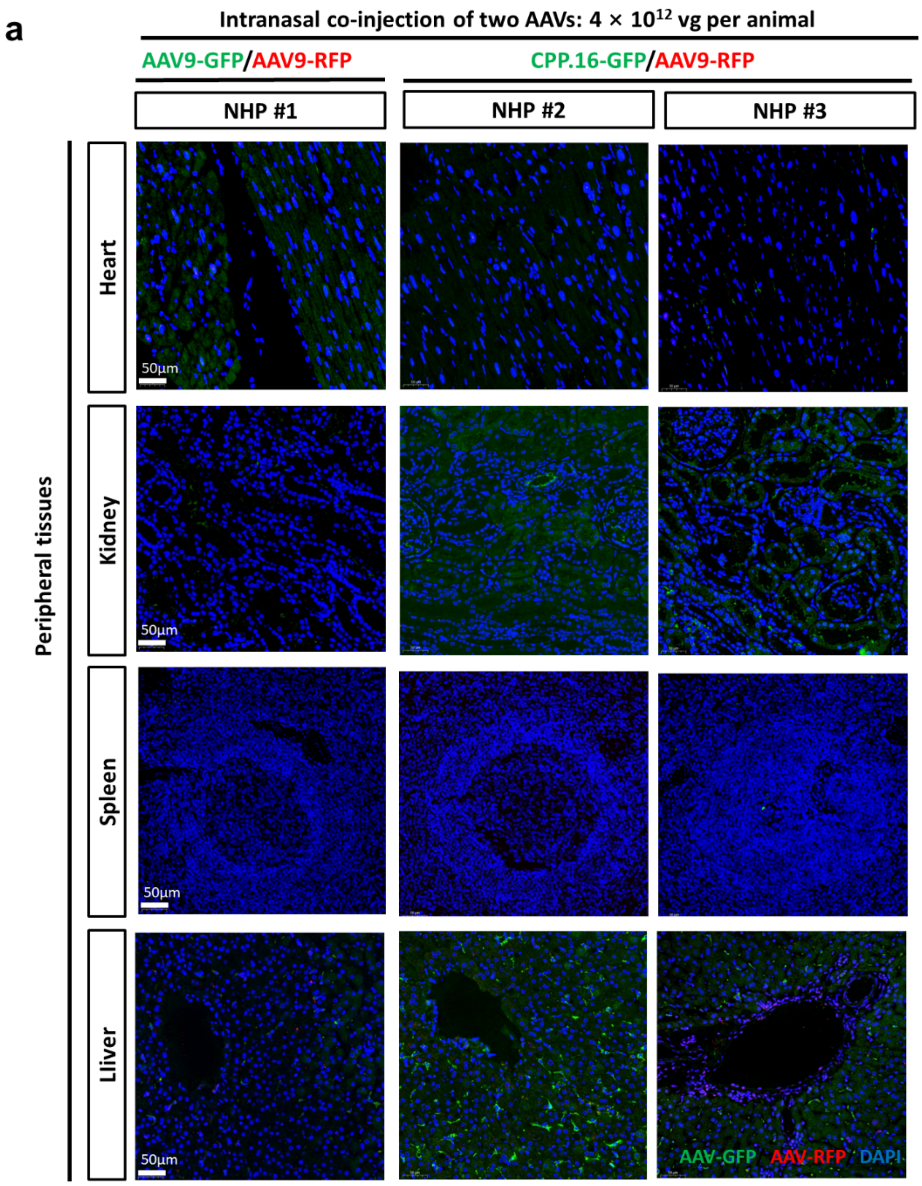


**Fig.S4.**related to Fig.3 **| AAV transduction in the heart, kidney, spleen and liver of the NHPs.**

Representative fluorescence images of GFP or RFP signals in the heart, kidney, spleen and liver of NHPs, 4 weeks after intranasal co-injection of total $4×10^{12}$ vg AAV9-GFP/AAV9-RFP or AAV.CPP.16-GFP/AAV9-RFP per animal. Scale bars: 50 µm.

#### Fig.S5. related to Fig.3

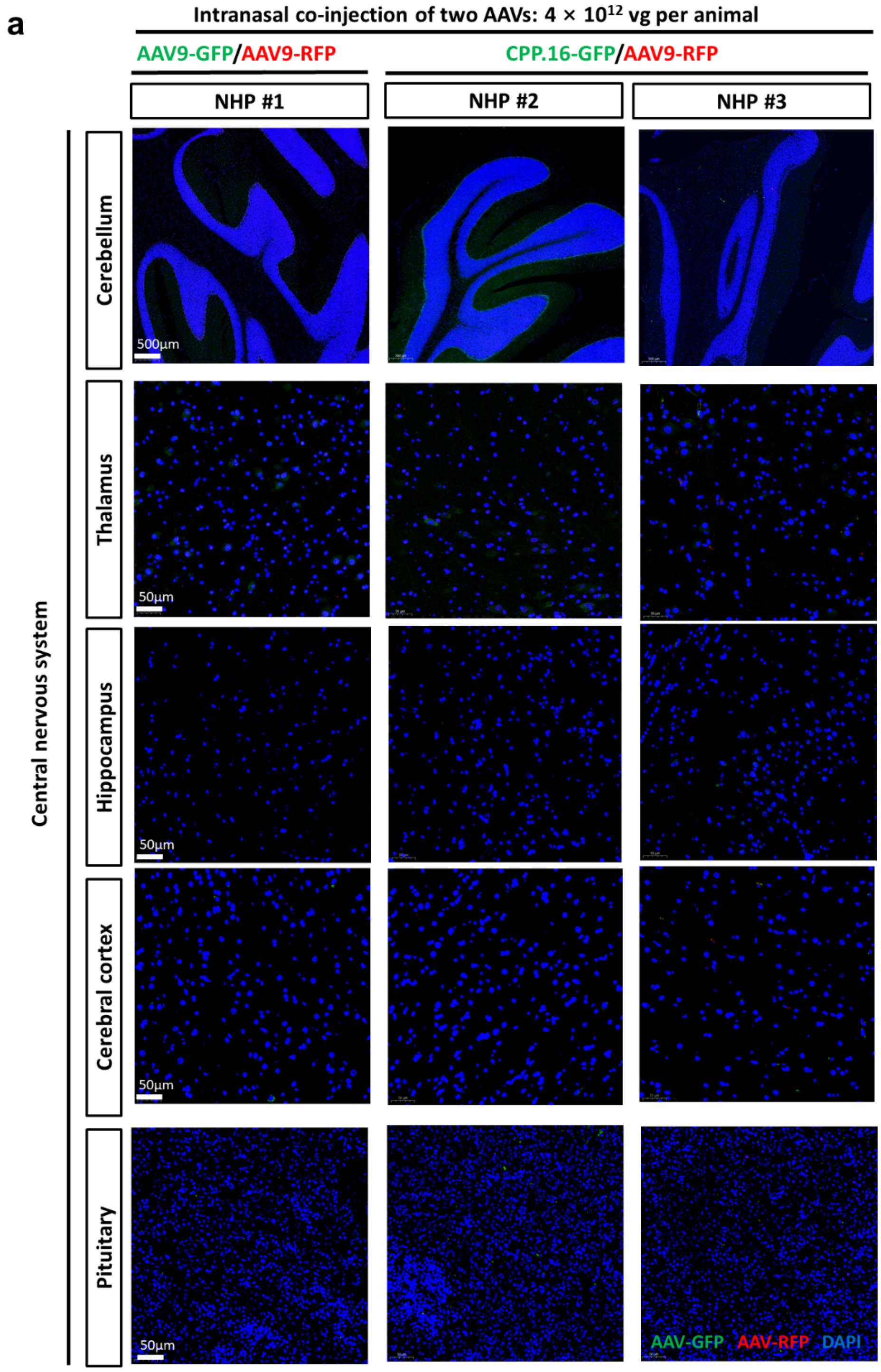


**Fig.S5.**related to Fig.3 **| No AAV.CPP. 16 transduction in the NHP brain after intranasal injection.**

**a**, Representative fluorescence images in the NHP brain, 4 weeks after intranasal co-injection of total $4{\times}10^{12}$ vg AAV9-GFP/AAV9-RFP or AAV.CPP.16-GFP/AAV9-RFP per animal. Scale bars: 500 μm for cerebellum sections and 50 μm for others.

#### Fig.S6. related to Fig.4

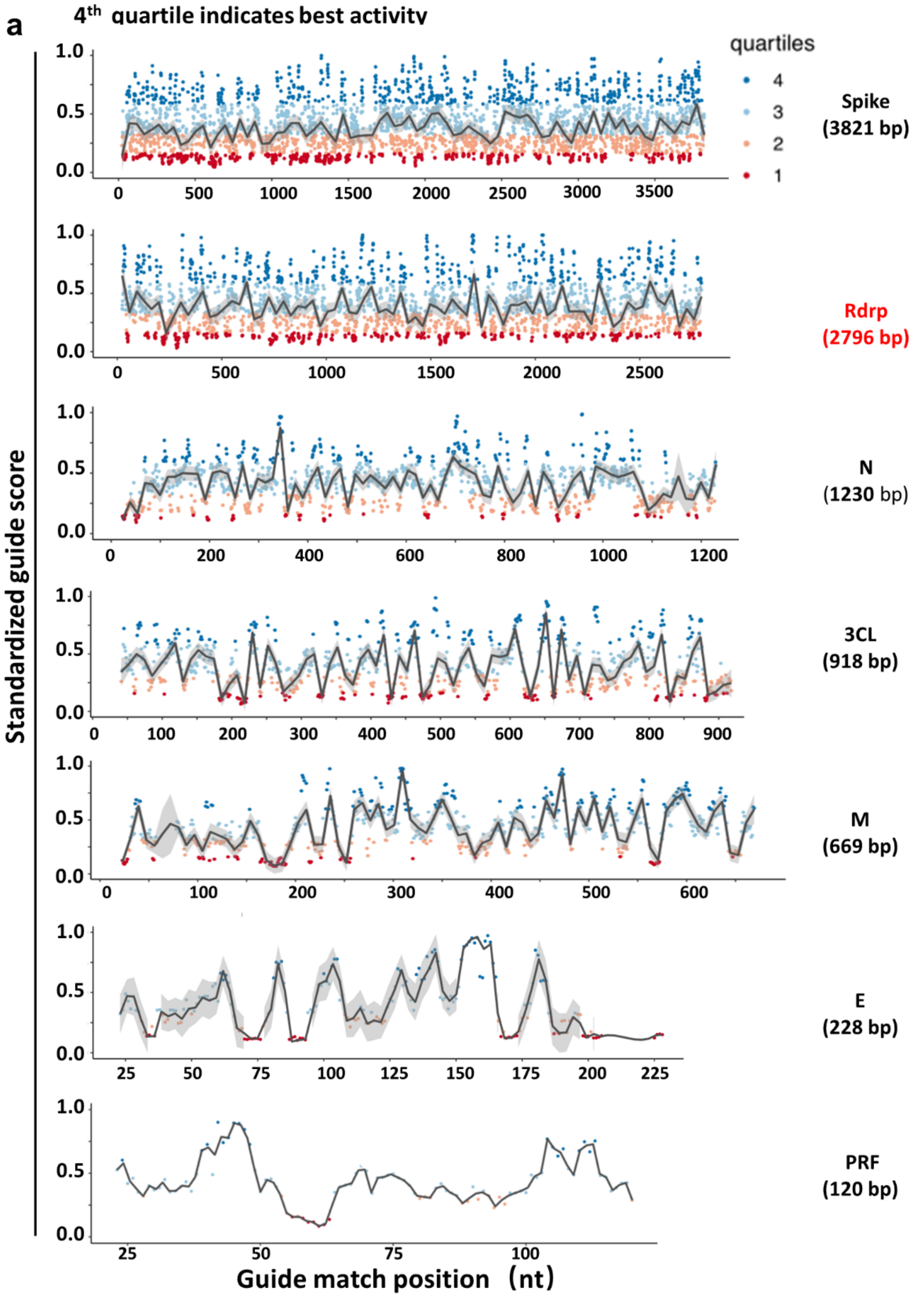


**Fig.S6.**related to Fig.4 **| Prediction of guide RNAs for conserved SARS-CoV-2 genes.**

**a**, Distribution and score of perfect-match guide RNAs for the conserved genes (spike, Rdrp, N, 3CL, M, E and PRF). Guide RNAs are separated into quartiles Q1-Q4 with Q4 containing guide RNAs with the best predicted knock-down efficiency.

#### Fig.S7. related to Fig.5

**a**

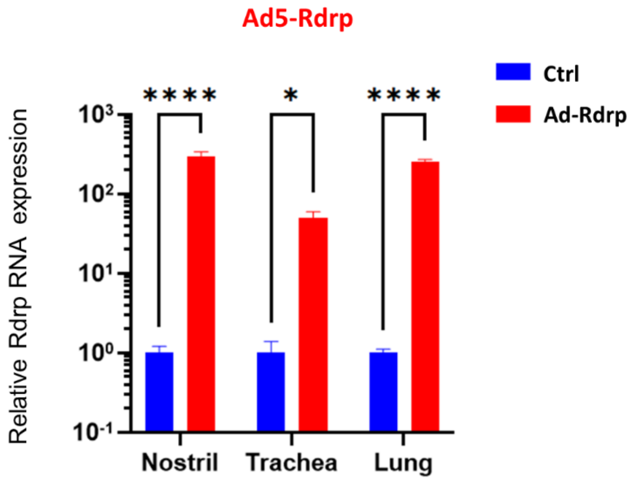


**Fig.S7.**related to Fig.5 **| Expression of Rdrp in mice by intranasal administration of Ad5-Rdrp.**

**a**, Quantification of Rdrp RNA expression in the nostril, trachea and lung 4 weeks after intranasal injection of Ad5-Rdrp in mice. n = 3 each group, mean ± s.e.m., two-way ANOVA with Sidak's post-test (*p < 0.05, ****p < 0.0001).
